# A conserved dimerization element is required for protein kinase activation by *trans*-autophosphorylation

**DOI:** 10.64898/2026.05.22.726961

**Authors:** Samuel Botterbusch, Vienna L. Huso, Kyler A. Weingartner, Gretar Tryggvason, Katherine W. Tripp, Rachel Green, Jennifer M. Kavran

## Abstract

*Trans-autophosphorylation* is the most common mode of protein kinase activation and involves two copies of the same kinase dimerizing so that one can phosphorylate the activation loop of the other. The diversity among structures of *trans*-autophosphorylation dimers supported the view that each kinase evolved a unique mode of recognition. We screened all human kinase crystal structures and identified an expanded set of dimers compatible with *trans-*autophosphorylation (655 dimers from 143 kinases). These dimers share no conserved structural arrangement, but 85% bury the same helix, αG, at the dimer interface. We validate αG-mediated dimerization by mutagenesis in kinases from each group of the kinome activated by *trans*-autophosphorylation. αG substitution impaired or abolished activation of full-length proteins, in cells, in every case. In purified kinase domains, αG substitution disrupted dimerization and autophosphorylation. These data establish that dimerization during *trans*-autophosphorylation is conserved and is mediated by a common structural element that, surprisingly, does not impose a specific arrangement of the two kinase domains relative to each other. αG is the least conserved element in the kinase fold, yet is required for activation across both the human kinome and other species, suggesting an ancestral function of the kinase fold.

**Significance Statement:** Protein kinases are the largest enzyme family in the human genome and common pharmaceutical targets. Most are activated by *trans*-autophosphorylation, during which two copies of the same kinase dimerize and one phosphorylates the activation loop of the other. How this is achieved remains poorly understood. We find dimerization during activation is conserved, but in an unexpected way. An unbiased structural screen of all human kinase crystal structures reveals kinases across the kinome bury the same helix, αG, at the dimer interface. αG-mediated dimerization extends to other species suggesting an ancestral function of the kinase fold. Substitution of αG disrupts activation of every kinase tested. Despite this conservation, αG-mediated dimers share no common arrangement, representing an unusual mode of protein-protein interaction.

## Introduction

The most widespread mechanism of protein kinase activation is *trans*-autophosphorylation, in which two copies of the same kinase dimerize, and one phosphorylates the activation loop (AL) of the other (1). How protein kinases dimerize during this process is assumed to differ, leaving the structural basis for kinase dimerization during activation restricted to studies of individual kinases. This assumption was well-founded. Inactive kinase domains sample a wide range of structural space, implying that each kinase is starting from a different point during activation (2). The relative arrangements of kinase domains in structures of *trans*-autophosphorylation dimers are not conserved (1, 3). The cellular inputs and regulatory domains that control dimerization differ across kinase families (2, 4).

Phosphorylation of the AL, which is achieved through multiple mechanisms including *trans*-autophosphorylation, is the central activating event for protein kinases. AL phosphorylation triggers a series of conformational changes throughout the kinase domain that stabilize the active conformation including re-ordering of the AL and repositioning components of the catalytic cleft to facilitate binding to substrates and ATP (2, 5, 6). The kinase domain consists of an N-lobe and C-lobe with the catalytic cleft formed between the lobes, and the AL extending from the C-lobe (***Figure 1A***). *Trans*-autophosphorylation is estimated to activate over half of the human kinome, including representatives from 7 of the 8 kinome groups (7). Yet structures of *trans*-autophosphorylation dimers have been characterized for less than 5% of the kinome (***Figure S1***). Individual structural studies have noted recurring features at the dimer interface, but whether any element is conserved and functionally required across the kinome has not been determined (8– 11).

**Figure 1.**
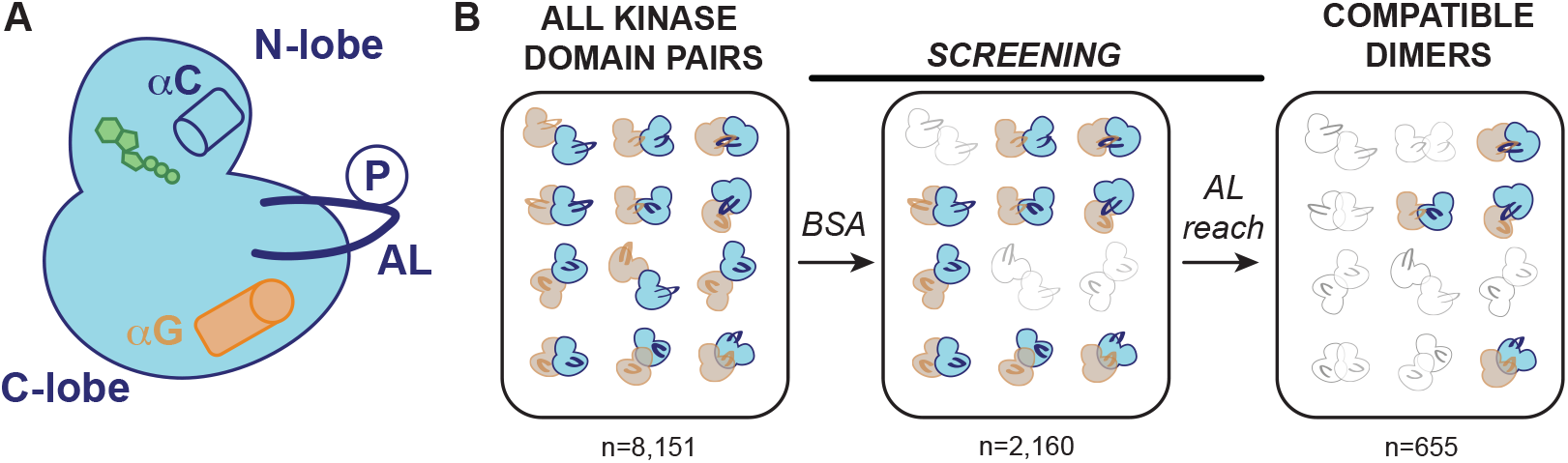
Screen to identify trans-autophosphorylation compatible dimers. (**A**) Cartoon schematic of an active, protein kinase domain showing the N-lobe, catalytic cleft (indicated by ATP in green), αC helix, the C-lobe, αG helix (orange), and a phosphorylated AL (indicated by a “P” in a circle). (B) Computational screen for *trans*-autophosphorylation-compatible dimers contains three steps: isolating all unique kinase-domain pairs, filtering for those that bury sufficient surface area, and AL reach modeling.

*Trans*-autophosphorylation has been difficult to characterize structurally. The interactions between the two kinase domains are weak and transient (8, 12, 13), the reaction proceeds through multiple steps, each with distinct geometric requirements, and the AL, which is the substrate of this process, undergoes large conformational changes. Prior identification of *trans*-autophosphorylation dimers required the AL to be resolved in one of two specific conformations (3, 8). The majority of crystal structures, however, fail to meet these conditions as the AL is frequently unresolved, suggesting that some trans-autophosphorylation compatible dimers have been either overlooked or underappreciated (14).

We developed a structural screen based on first principles to overcome these limitations and address whether or how dimerization is conserved during activation. Rather than requiring a specific AL conformation, this screen identifies dimers compatible with *trans*-autophosphorylation, specifically that the AL could be modeled to reach the opposing catalytic cleft. Using all available human protein kinase crystal structures, 655 homodimers of 143 kinases, spanning the kinome, were identified by this screen. These dimers share no conserved dimer arrangement, yet 85% of them bury the same helix at the interface – αG, a small non-catalytic helix on the C-lobe (***Figure 1A***). The functional requirement of αG in *trans*-autophosphorylation was tested for a subset of kinases from each kinome group activated by this mechanism. For each kinase tested, substitution of αG impaired or abolished activation of full-length proteins in cells. For purified, isolated kinase domains substitutions of αG residues did not alter the kinase domain fold but did block dimerization. These data establish that dimerization during *trans*-autophosphorylation is conserved kinome-wide and is mediated by the same helix. Unexpectedly, this conserved dimerization element does not impose a conserved dimer arrangement.

## Results

### A structural screen identifies trans-autophosphorylation compatible dimers across the kinome

We developed a structure-based screen to identify *trans*-autophosphorylation compatible dimers of human protein kinases (***Figure 1B***). We analyzed one representative of each unique crystal form available in the Protein Data Bank to create a non-redundant data set (***Supporting Information 1***). We then generated all possible kinase domain pairs, mediated by either contacts in the asymmetric unit or through crystal symmetry, and identified those that satisfied two criteria. First, candidate dimers must bury at least 500Å^2^ of surface area per monomer, a threshold for biologically relevant contacts (15, 16). Second, the AL of at least one kinase must be capable of reaching into the catalytic cleft of the other, assessed by modeling the shortest sterically allowed path. This approach identified 655 homodimers of 143 kinases, including representatives from all kinome groups (***Figure S1, Supporting Information 2***). This screen recovered 20 of the 22 documented *trans*-autophosphorylation crystal structures available at the time; two dimers were excluded because of steric clashes between sidechains that prevented AL path modeling.

### Trans-autophosphorylation dimers do not share a conserved arrangement

No dimer arrangement is used consistently across the kinome, as shown by pairwise superpositions of all 655 dimers onto each other (***Figure 2A***). This lack of conservation was also true within individual kinome groups (***Figure 2B***). Three small clusters of similar arrangements were identified, but they corresponded to dimers of single kinases (IRAK4 or MAPKAPK2) or closely related family members (DAPK family), reflecting features specific to a given kinase rather than the kinome. The latter corresponds to a known, autoinhibitory homodimer (17). For individual kinases with multiple dimers, arrangements were generally heterogeneous (***Figure S2***).

**Figure 2.**
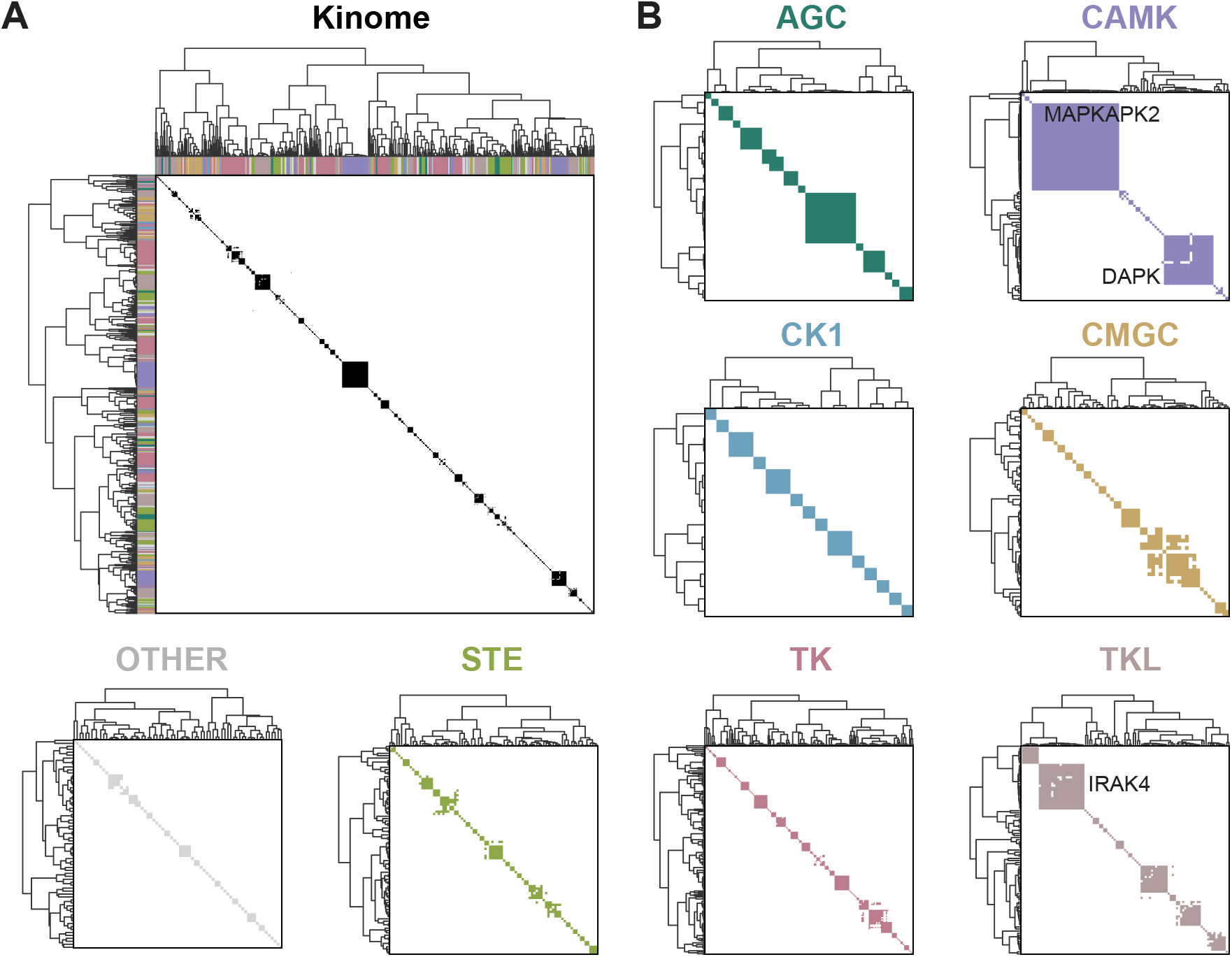
Trans-autophosphorylation compatible dimers are structurally diverse. (**A**) Pairwise superpositions of all *trans*-autophosphorylation dimers clustered by similarity based on root mean squared deviation (RMSD); white indicates not conserved ( RMSDs of ≥3Å) and black conserved (RMSDs ≤ 3Å). Colors along each axis indicate the kinome group of each kinase as colored in panel B. (**B)** RMSD calculations as in (A), calculated within individual kinase groups. The three small clusters of similar arrangements are identified by name. Number of dimers analyzed: 625 kinome, 29 AGC, 89 CAMK, 17 CK1, 56 CMGC, 71 Other, 90 STE, 100 TK, and 100 TKL. 30 structures were excluded from this analysis because residues used in the superpositions were disordered.

### αG is the most frequently buried element at the dimer interface

Despite this structural heterogeneity, the same two regions were consistently buried at the interface for 85% of the dimers, the AL and αG-helix (***Figure 3A, B***). The screen captured αG-mediated dimers for 130 out of 143 kinases (***Figure 3C***). In these αG-mediated dimers, the αG of one kinase preferentially contacted the αG and AL of its partner (***Figure S3***). When analyzed by kinome group, the AL and αG remained the most frequently enriched elements at the dimer interface for seven out of eight groups (***Figure 3D, S4***). The one group that was the exception, CK1, is not activated by *trans-*autophosphorylation (18). Each group also had a distinct pattern of additional, preferentially buried residues, suggesting dimerization preferences evolved within each group. Preferential use of αG is only found in the subset of dimers (655) that satisfy both screening criteria and not for subsets that satisfy only one or the other criteria (***Figure S5***).

**Figure 3.**
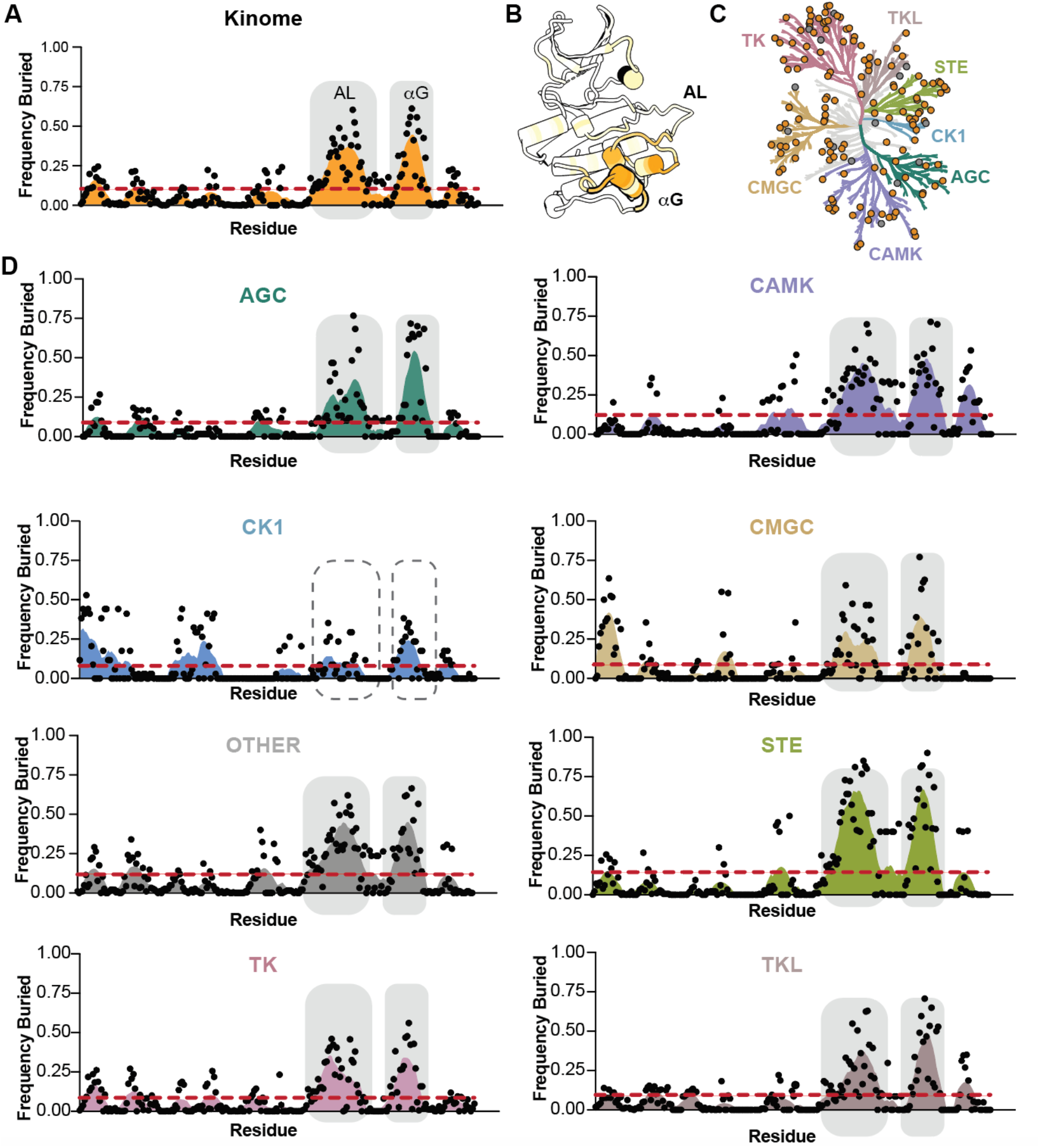
αG is the most frequently buried element at the dimer interface. (A) Dimer footprint plot showing the frequency with which residues in conserved secondary structure elements are buried at the dimer interface across all *trans*-autophosphorylation compatible dimers. Dashed red line indicates the mean frequency; orange area is a smoothed fit for visualization; gray boxes highlight the two most frequently buried regions. (B) Cartoon representation of phosphorylated MST1 kinase domain (3COM) colored by frequency buried (dark orange, high; light orange, low). (C) Human kinome in which each dot represents a kinase with a *trans*-autophosphorylation compatible dimer. Orange dots indicate αG-mediated dimers, gray dots indicate not αG-mediated dimers. (D) Dimer footprint plots as in *(A*) calculated by kinome group. Number of dimers analyzed: 655 kinome, 30 AGC, 91 CAMK, 17 CK1, 59 CMGC, 91 OTHER, 90 STE group, 174 TK, and 104 TKL.

If αG-mediated dimerization is a general feature of *trans*-autophosphorylation then it should be present in dimers identified by other methods, in other species, and in solution. This is the case. Every documented structure of a human *trans*-autophosphorylation dimer has αG at the interface. Three different methods were used to identify these dimers (AL exchange, geometric positioning of catalytic elements, or the structural screen used here), and all determined αG-mediated dimers (***Figure 4A***) (3, 8, 10, 12, 19–32). These structures include MEKK2, which was published during preparation of this manuscript (9). αG*-*mediated *trans*-autophosphorylation dimers have been observed across eukaryotes, including mouse, salmon, and yeast, and even in the eukaryotic-like kinases found in mycobacteria (***Figure 4B***) (33–36). αG-mediated dimers have been captured by cryoEM and analytical ultracentrifugation (AUC) confirming these contacts form in solution (10, 33). αG’s role in dimerization and AL autophosphorylation has been validated for nine kinases but whether this role was conserved kinome wide was not investigated (***Table 1***) (3, 10, 22, 23, 25, 27, 37–39).

**Table 1.** Evidence for αG in dimerization and activation across the kinome.

| Group | Kinase | Cell based autophosphorylation | In vitro dimerization and/or autophosphorylation | Reference |
| --- | --- | --- | --- | --- |
| AGC | PKD1 | n.d.<sup>^</sup> | Y2888A* | (38) |
|  | PKD1 | n.d. | Y2888E* | (38) |
|  | NDR1 | E325A, T326A, Q328A, K332A, M335A, N336A | n.d. | Present work |
| CAMK | CHK2 | R431A, T432A, Q433A, V434A, S435A, K437A, D438A, T441A, S442A | R431A, T432A, Q433A, V434A, S435A, K437A, D438A, T441A, S442A | Present work |
| CMGC | JNK2 | n.d. | I231A | (39) |
|  | JNK2 | I231A | n.d. | Present work |
|  | JNK2 | T228A, D229A, H230A, I231A, E239A | n.d. | Present work |
| STE | PAK1 | L470A/D | n.d. | (25) |
|  | PAK1 | L473A | n.d. | (25) |
|  | PAK2 | n.d. | L499Q | (37) |
|  | PAK2 | n.d. | L452Q | (37) |
|  | MEKK2 | M567D, I570D, F571A | n.d. | (9) |
|  | MEKK2 | M567D, I570D, F571D | n.d. | (9) |
|  | MST2 | Y221, F231, M232, T235, N236, P237, P238 | Y221, F231, M232, T235, N236, P237, P238 | (10) |
|  | MST2 | K218, D223, I224, H225, P226, M227, R228 | K218, D223, I224, H225, P226, M227, R228 | (10) |
|  | MST4 | M225E, L228E, F229A | n.d. | (27) |
| TK | LCK | T445V | n.d. | (3) |
|  | LCK | N446A | n.d. | (3) |
|  | FES | Q768A, F772A, K775A | n.d. | Present work |
|  | FES | F772A | n.d. | Present work |
|  | IGF1R | N1218A | n.d. | Present work |
|  | IGF1R | R1223E | n.d. | Present work |
| TKL | IRAK4 | n.d. | E401K | (22) |
|  | IRAK4 | Q394A, L395A, D398A, E401A, E404A, D405A, E406A | n.d. | Present work |
|  | IRAK4 | E401K | n.d. | Present work |
|  | ZAK | E210K, L212A, W216A, E220K, K221E | n.d. | Present work |
|  | ZAK | K221E | n.d. | Present work |
| OTHER | AuroraA | n.d. | C290A | (23) |

\* data only available for autophosphorylation
^ not determined (n.d.)

**Figure 4.**
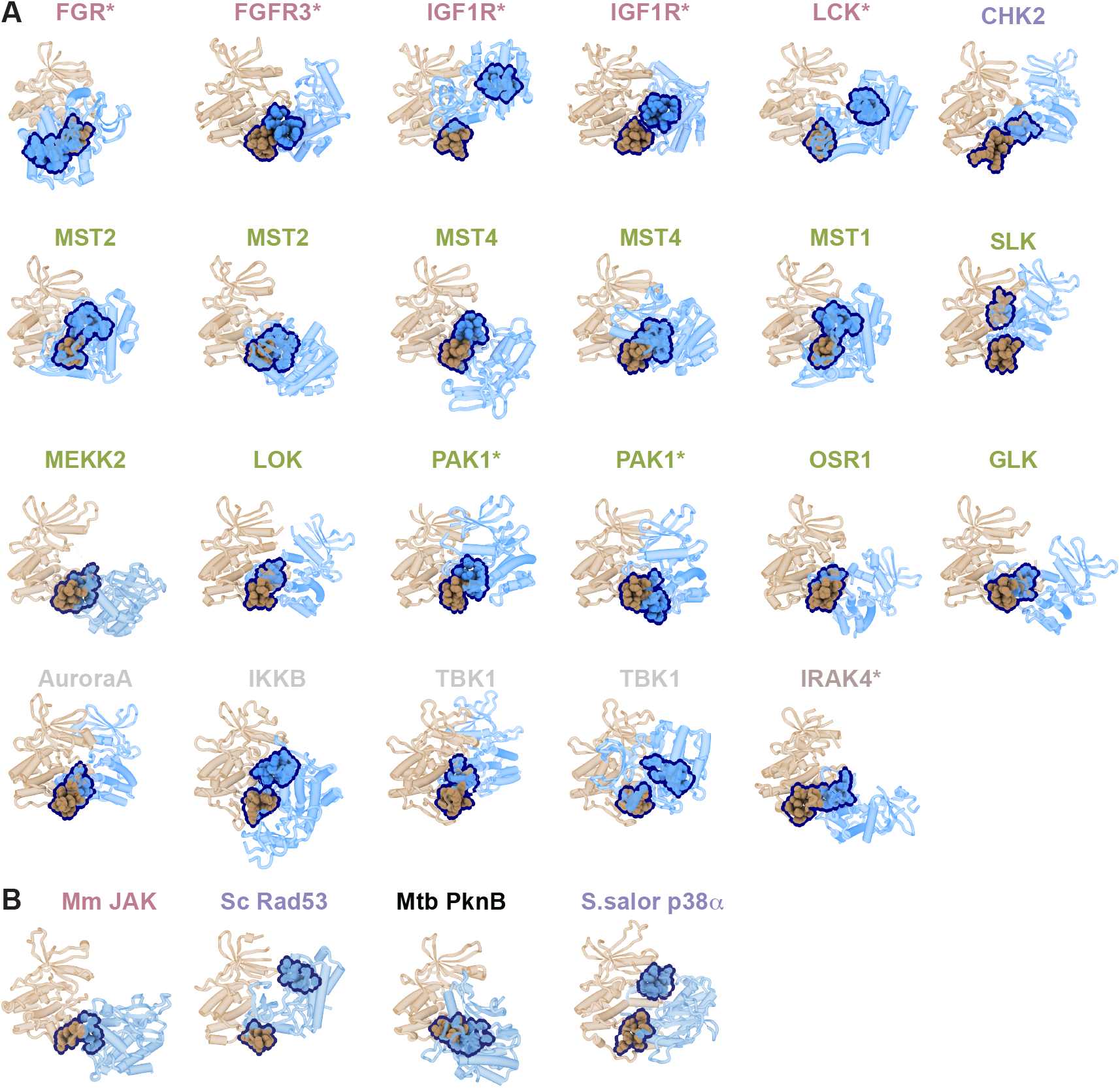
The G-helix mediates dimerization in all documented trans-autophosphorylation structures. (**A**) Cartoon representations of 23 *trans*-autophosphorylation structures from five kinome groups (AGC in teal; CAMK in purple, Other in gray, STE in light green, TK in light pink and TKL in light brown). In each dimer, one kinase domain is shown in tan and the other blue. αG residues are displayed in spheres. Dimers that represent the transition state are indicated by an asterisk. PDBIDs: *Row 1 --* 7UY0, 6PNX, 3D94, 3LVP, 2PLO, 2YCF; *Row 2 --* 8A66, 4LG4, 4FZA, 3GGF, 3COM, 2J51; *Row 3—*9P6A, 2J7T, 3Q4Z, 4ZJI, 3DAK, 5J5T ; *Row 4 --* 4C3P, 4KIK, 4EUT, 4EUU, 4U97. (**B**) Cartoon representation of *trans*-autophosphorylation dimers from other species including *Mus musculus* (8EWY), *Saccharomyces cerevisiae* (4PDP), *Mycobacteria tuberculosis* (3F69), and *Salmo salor* (3OHT). Structures displayed and labelled as in A, and the eukaryotic-like S/T protein kinase is labelled in black.

### αG is functionally required for activation

We tested the functional requirement for αG by monitoring the effects of substitutions of surface-exposed αG residues (αG*) for a set of kinases from groups for which the role of αG had not been fully established: NDR1 (AGC group), CHK2 (CAMK group), JNK2 (CMGC group), FES and IGF1R (TK group), and IRAK4 and ZAK (TKL group). Full-length kinases were transiently expressed in cells, triggering their activation, and AL phosphorylation compared between αG* variants and either wild-type (WT) or kinase-inactive (KI) controls using either phosphospecific Western blots (IRAK4, CHK2, and JNK2) or pan-phosphotyrosine blots after immunoprecipitation (FES). For each kinase, AL autophosphorylation was reduced in αG* variants compared to wild-type (***Figure 5A-C***). Single-amino acid substitutions were sufficient to block activation for JNK2, IRAK4, and FES (***Figure S6A***,***B***,***C***).

**Figure 5.**
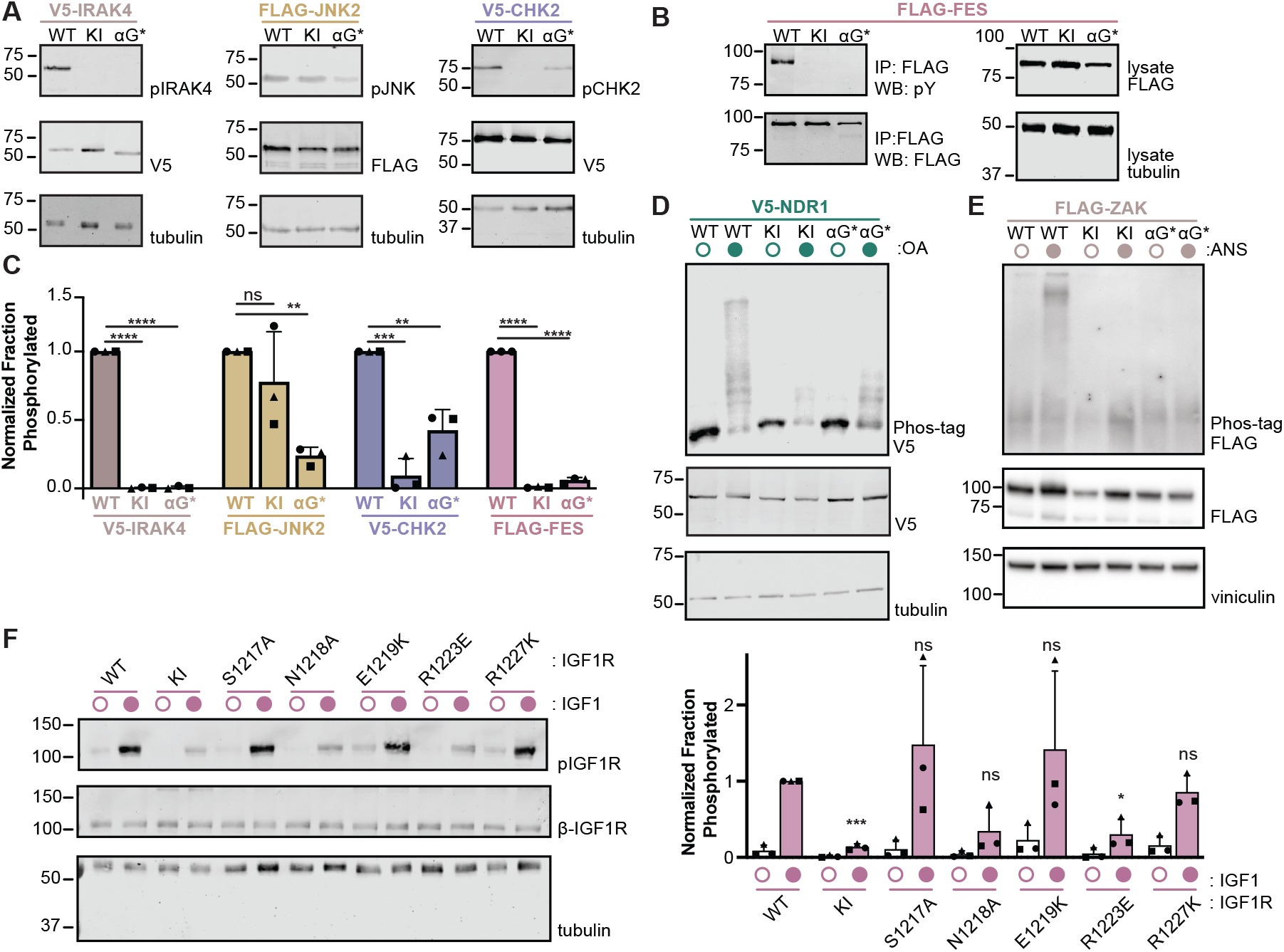
αG is required for activation in cells. Representative Western blots of HEK293 lysates transiently transfected with IRAK4, JNK2, CHK2 (A) and, for FES (B), lysates followed by immunoprecipitations, probed with indicated antibodies. (C) Bar graph of the mean with standard deviation of normalized, fraction phosphorylated protein. Biological replicates are indicated by different shaped points. Significance determined by one-way ANOVA followed by Dunnett’s test. (D, E) Lysates expressing indicated variants of either NDR1 (*D*) or ZAK (*E*) resolved on either Phos-tag (*top*) or standard SDS-PAGE Western blots (*bottom*) probed with indicated antibodies. (F) (*Left*) Representative Western blots of HEK293 lysates transiently transfected with indicated IGF1R variants and probed with indicated antibodies. (*Right*) Bar graph of the mean with standard deviation of normalized, fraction phosphorylated IGF1R for three biological replicates, and plotted as in *C*. Significance determined by one-sample t-test between samples with IGF1. P values used: ****<0.0001; *** <0.001, ** < 0.01; *<0.05, and ns >0.05. ANS stands for anisomycin; OA for okadaic acid.

Activation of ZAK and NDR1 kinases was monitored by Phos-tag gel, for which protein migration reports on overall, rather than site-specific, phosphorylation because phospho-specific antibodies for the AL were not available. NDR1 is activated in a two-step process; phosphorylation of its hydrophobic motif by an upstream kinase followed by autophosphorylation of its AL (40). Prior to activation, all NDR1 variants migrated as a single species. Following addition of okadaic acid, wild-type NDR1 super-shifted and migrated as multiple species. αG* variants shifted only partially, similar to kinase-inactive controls, consistent with phosphorylation of the hydrophobic motif but no phosphorylation of the AL (***Figure 5D***). ZAK is activated by ribosome collisions resulting in autophosphorylation of both its AL and multiple sites on its C-terminal tail (41–44). Anisomycin treatment, which triggers ribosome collisions, resulted in a super-shift of wild-type ZAK but no change in the migration of either αG* variant or the kinase-inactive control, indicating inhibition of autophosphorylation (***Figure 5E***). Single amino-acid substitutions of αG in ZAK revealed graded, site-specific effects (***Figure S6D***).

An αG* variant of IGF1R had impaired processing of the pro-receptor, so we screened five single-residue substitutions of surface exposed, αG residues instead. Activation was triggered by addition of ligand, IGF1, and AL phosphorylation monitored with phospho-specific Western blot. Two different single-substitutions of αG, N1218A and R1223E, reduced AL phosphorylation compared to wild-type, while three other substitutions, S1217A, E1219K, and R1227K, each had no effect (***Figure 5F***). The impairment of autophosphorylation of the AL is therefore a consequence of site-specific changes.

### αG is required for kinase domain dimerization

The cell-based data establish that αG is required for activation but do not report on whether it is needed for dimerization *per se*. Purified wild-type and αG* CHK2 kinase domains have identical secondary structure, as judged by far-UV circular dichroism (CD), confirming αG substitutions do not disrupt the kinase fold (***Figure 6A***). Wild-type CHK2 sedimented in solution as a mixture of species in a concentration-dependent fashion, as monitored by analytical ultracentrifugation (AUC), indicative of a monomer to dimer equilibrium. αG* CHK2 kinase domain, in contrast, sedimented as a single species across the same concentration range, consistent with monomeric protein (***Figure 6B***). αG* CHK2 kinase domain in solution also had impaired autophosphorylation compared to wild type (***Figure 6C***). The activation defects observed in cells can be directly attributed to disruption of αG-mediated dimerization.

**Figure 6.**
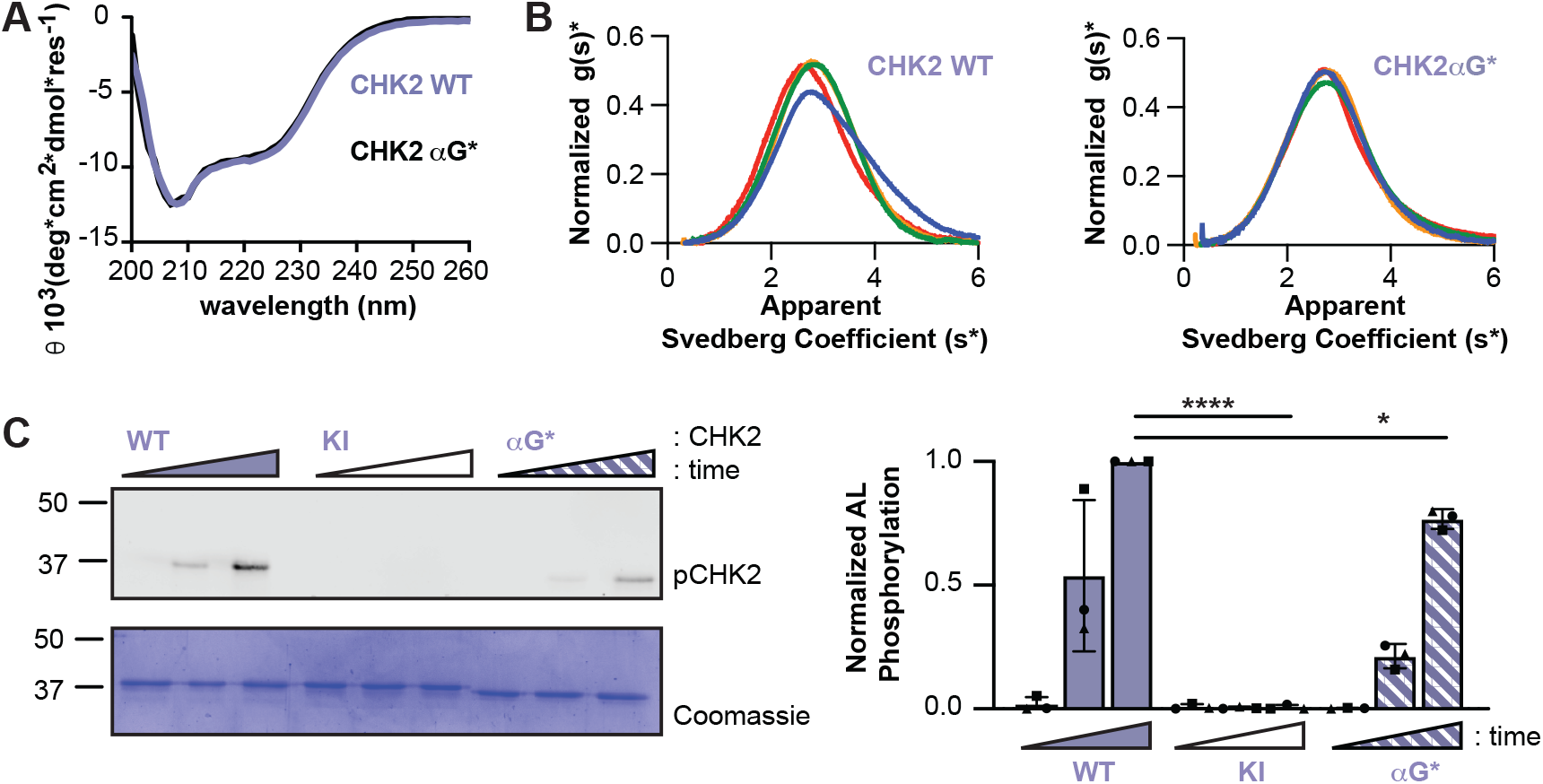
αG substitutions disrupt CHK2 kinase domain dimerization and autophosphorylation. (**A**) Overlay of far-UV CD spectra for CHK2 wild-type kinase domain (light purple) and CHK2 αG* (black). (**B**) Sedimentation coefficient distributions of CHK2 kinase domain wild type (*left*) or αG* variant (*right*) displayed as g(s*) plots. Curve colors correspond to protein concentration (0.3 mg/mL in red, 0.6 mg/mL in orange, 0.9 mg/mL in green, 1.2 mg/mL in blue). (**C**) Purified CHK2 kinase domain variants incubated with ATP and samples taken at three time points. (*Left*) Representative blots analyzed either by phospho-specific antibody or stained with Coomassie. (*Right*) Bar graph of the mean fraction phosphorylated from three experiments, and error bars indicate the standard deviation. Statistical significance calculated by one-way ANOVA of last time points (****p <0.0001 and *p<0.05).

## Discussion

αG mediates dimerization during *trans*-autophosphorylation across the kinome. 85% of dimers in our data are αG-mediated (***Figure 3***). Substitutions of αG disrupt AL autophosphorylation for every kinase tested, both in full-length proteins in cells and purified kinase-domains, establishing αG as functionally required (***Figures 5***,***6, S6, Table 1***). Single-amino acid substitutions are sufficient to block activation in a site-specific manner for multiple kinases (***Figure 5F, S6***). For NDR1, which requires both upstream activation and *trans*-autophosphorylation, only the latter is controlled by αG (***Figure 5D***). αG substitution blocks dimerization of isolated kinase-domains and inhibits autophosphorylation without disrupting the kinase fold for two kinases, shown here for CHK2 (CAMK group) (***Figure 6)*** and in our prior work for MST2 (STE group) (10). This function for αG is unexpected; while often noted to mediate protein-protein interactions (11), it is not part of either the regulatory or catalytic spines, is distal from the catalytic cleft and αC-helix, and has no direct catalytic function.

αG-mediated dimers are unusual because while αG is found at the dimer interface across the kinome, the residues it buries, the residues it contacts on the partner kinase, and the orientation of the two kinase domains within the dimer vary. The buried element is conserved but nothing else about the interaction is, regardless of whether comparing dimers across the kinome, within individual groups, or between structures of the same kinase (***Figures 2, S2***). This extends even to dimers that capture the transition-state, which has the most precise geometric constraints; two transition-state dimers have been determined for each of IGF1R and PAK1, and in both cases the dimer arrangements are unique (***Figure 4A***). The absence of a conserved arrangement among transition-state dimers indicates that dimerization provides proximity and that productive engagement of the AL as a substrate is a separate event. Upstream events that trigger kinase activation typically increase the local concentration of the kinase-domain, thereby promoting formation of αG-mediated dimers (45). In the absence of these cues, αG-mediated dimerization is likely prevented by either autoinhibitory domains and/or low concentration.

αG-mediated dimerization extends beyond *trans*-autophosphorylation to *trans*-phosphorylation, another mode of activation where the AL of one kinase is phosphorylated by a different, upstream kinase, and is often found in signaling cascades. Seven of the eight experimentally determined *trans*-phosphorylation structures are αG-mediated and have heterogenous dimer arrangements (***Figure S7***) (46–53). For six *trans*-phosphorylation pairs, αG has been experimentally validated in activation (9, 49, 50, 54–57). The shared role of αG in both *trans*-autophosphorylation and *trans*-phosphorylation points to a common structural basis for dimerization-driven kinase activation. Beyond activation, αG functions as a multifunctional docking site, mediating substrate recognition, scaffolding, inhibitor binding, autoinhibition, and allosteric activation on an individual kinase basis (11, 58). These functions differ between kinases (for example CK1 group uses αG for thermal regulation and not homodimerization), unique and separable for a single kinase (for example EGFR (TK group) uses αG both in its allosteric, asymmetric dimer and inhibition by MIG6) (18, 59, 60). This multifunctionality suggests that distinct αG-mediated interactions are competitive and may impose a hierarchy to ensure fidelity of signal transduction. The physical properties of αG explain both this functional versatility and diversity of dimer arrangements it mediates. αG is part of the GHI helical expansion that defines the eukaryotic protein kinase fold (61). It is in every kinase but is the least conserved element in the fold, has a variable position relative to the catalytic core, and has no direct catalytic role (62).

αG mediates dimerization during *trans*-autophosphorylation across the kinome and the evolutionary tree. Our findings predict αG as a site for inactivating mutations and new target for therapeutic intervention. To paraphrase Tolstoy, all *trans*-autophosphorylation dimers are alike in αG-mediated dimerization; each dimer is arranged in its own way. The least conserved structural element in the protein kinase fold is its most versatile.

## Materials and Methods

### Identification of trans-autophosphorylation compatible dimers

Every available human protein kinase structure was downloaded from KLIFS on July 17, 2023 (63). Those structures were pruned to contain a single entry for each unique crystal form based on protein sequence, phosphorylation state, space group, and unit cell dimensions (linear cell variation ≤2%). If multiple structures were available for the same crystal form, then the highest resolution one was kept (64) (***Supporting Information 1***). Any non-kinase-domain atoms were then removed and residues renumbered according to a universal numbering system (62). For each structure, every kinase domain pair was generated by applying crystallographic symmetry to the asymmetric unit in PyMOL (The PyMOL Molecular Graphics System, Version 3.0 Schrödinger, LLC) (***Supporting Information 2***).

Pairs were classified as *trans*-autophosphorylation compatible dimers if they satisfied two criteria. Buried surface area was calculated using areaimol in CCP4 and a ≥500 Å^2^/monomer cutoff applied (15, 16, 65, 66). AL modeling was performed in two steps. Pairs were first triaged using straight-line distance calculation between the Cαs of F1339 (DFG motif, universal numbering) of one kinase to D1013 (HRD motif) of its partner and back to P1915 (APE motif). If this distance was shorter than the length of a fully extended AL (number of residues x 3.4Å), the pair was included for further modeling. The shortest path for the ALs through unoccupied space were then modelled. AL residues (F1339 - P1915) were removed from the model. The remaining structure was placed in a 1Å^3^ grid with boundaries twice the dimensions of the kinase pair. The AL was modelled as a tube with a radius of 3.5Å and restrained to not approach within 1.6Å of the center of any non-hydrogen atom (67). The shortest grid path through unoccupied space was calculated using Dijkstra’s algorithm in two steps. The path started from the nearest unoccupied point from Cα-D1013 and proceeding to each grid point continuing to either Cα-F1339 or Cα-P1915. These path lengths were then combined and smoothed to the shortest path length using a dampened elastic string model (68)(***Equation 1***):

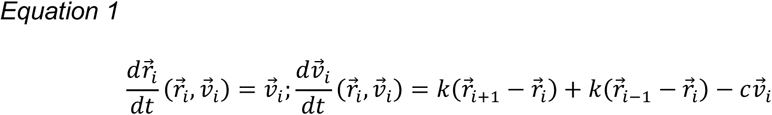

Where k is the spring constant (0.1 dt^-2^), c is the drag constant (0.4 dt^-1^), and r and v are position and velocity vectors, respectively. The path was allowed to relax for 500 iterations or until convergence (no shortening for 10 consecutive iterations).

### Structural Analysis

Buried residues were identified using areaimol in CCP4 (65). RMSD calculations were performed with ProDy (69) and were calculated using 72 Cα atoms on the C-lobe of the kinase domain. The RMSD heatmaps were generated by first calculating all permutations of pairwise RMSDs and then using hierarchical clustering of the lowest RMSD in scikit-learn with a 3Å RMSD threshold for cluster membership(70).

### Cloning

DNA encoding the kinase domains of human CHK2 was obtained from the Human Kinase Domain Constructs Kit, a gift from John Chodera (Addgene #1000000094). Full-length human CHK2 (Addgene #41901, gift from Stephen Elledge) and NDR1 (Addgene #37023, gift from Yutaka Hata) were cloned into pcDNA3.1(+) (Invitrogen) with an N-terminal V5 epitope tag. DNA encoding either full-length, human FES or JNK2 (GenScript Clones ID OHu28580D and OHu19851D, respectively) were cloned into pcDNA3.1(+) with a C-terminal FLAG epitope tag. DNA encoding full-length human ZAK with an N-terminal 3xFLAG tag under a partial CMV promoter (pFN24K) was cloned previously (41). DNA encoding full-length IGF1R in a modified pSGHV0 was cloned previously(71).

Standard molecular biology techniques were used to generate site-specific substitutions. αG* mutations included alanine substitutions of any αG-residue with more than 30% solvent accessible surface area, as calculated in PyMOL. For IGF1R, FES, and ZAK this cluster was broken into single substitutions, including charge-swaps, as indicated. Single-site substitutions of IRAK4 and JNK2 were based on those previously reported for purified systems(22, 39). A full list of variants used is in ***Table S1***.

### Kinase domain expression and purification

Plasmids encoding, separately, a His-tagged CHK2 kinase domain and MBP-tagged lambda phosphatase was transformed into *E*.*coli* Rosetta(DE3) (Novagen). Cells were grown overnight at 25°C in Terrific Broth Autoinduction Media (Grisp Biosciences), as described (72). CHK2 was purified using nickel-charged Profinity immobilized metal affinity chromatography (IMAC) resin (BioRad), followed by gel filtration chromatography. Protein was concentrated to 12 mg/mL in 10mM HEPES pH 8.0, 150mM sodium chloride, 500µM TCEP (tris(2-carboxyethyl)phosphine) and flash frozen.

### Tissue Culture

HEK293T cells (ATCC) were cultured in DMEM:F12 medium (Gibco) supplemented with 5% fetal bovine serum (VWR) and 2mM glutamine (Gibco). HEK293T ZAK knockout cells were maintained in Dulbecco’s modified eagle medium (DMEM; Thermo) supplemented with 10% FBS (Thermo Fisher)(41). Cells were grown at 37°C with 5% CO_2_.

### In-cell activity assays

For all kinases except ZAK, the following protocol was used. HEK293T cells were seeded at 0.3×10^6^ cells/well, transfected with polyethyleneimine with indicated plasmids (73) or for CHK2 FuGene HD (Promega). Cells were lysed 48 hours post-transfection using RIPA supplemented with phenylmethanesulfonyl fluoride (PMSF), protease inhibitor cocktail (Sigma), sodium vanadate, sodium fluoride, sodium pyrophosphate, ß-glycerophosphate, EDTA, and Benzonase (Millipore Sigma). Clarified lysates were normalized using bicinchoninic (BCA) assay (Pierce) and resolved by Western blot using indicated antibodies ***Table S2***. Samples were loaded to ensure signal was within linear range of antibody and detection. Blots were scanned with Odyssey (LiCOR Bio), and band intensities quantified in ImageJ (74). Fraction phosphorylated was calculated as the ratio of phosphorylated to total band intensity for each protein and then normalized to the wild-type within each replicate. 3 biological replicates were obtained for each in-cell assay.

The above protocol was adapted as follows. For NDR1, cells were treated with either 1 μM okadaic acid (ThermoFisher) or vehicle one hour prior to lysis. Lysates were resolved on either standard or Phos-tag SDS-PAGE (7% 29:1 acrylamide, 21µM Phos-tag acrylamide (FujiFilm), and 42µM manganese chloride) followed by Western blot. For FES, protein was immunoprecipitated from lysates using an anti-FLAG antibody. For IGF1R, cells were seeded at 1×10^6^ cells/well, then at 16 hours post-transfection serum-starved for 3 hours in the presence of 1mg/mL BSA and then incubated with either 20nM IGF1 (PeproTech) or vehicle for 30 minutes before lysis.

ZAK in cell assays followed previously published protocols(41). Briefly, ZAK knockout cells were transfected with indicated plasmids using Lipofectamine 3000 (Thermo). After 24 hours, media was changed and anisomycin (Sigma) added to the media 15 minutes prior to lysis. Clarified lysates were resolved by Western blot using indicated antibodies (***Table S2***) on either standard reducing SDS-PAGE or Phos-tag SDS-PAGE (8% 29:1 acrylamide, 10.7 µM Phos-tag acrylamide (FujiFilm) supplemented with 21.3 µM manganese chloride). Blots were scanned with Bio-Rad ChemiDoc imaging system (SuperSignal West Pico PLUS and West Femto Maximum reagents, Thermo).

### Analytical Ultracentrifugation

All proteins were dialyzed for 24 hours in their indicated storage buffer and then diluted to 1.2, 0.9, 0.6, and 0.3 mg/mL. Samples and reference were loaded into ultracentrifuge cells assembled with SedVel60K 1.2cm centerpieces, and menisci matched by spinning at 50,000 rpm in a Beckman XL-I. Following meniscus-matching, all cells were rocked and inverted to homogenize any generated concentration gradient. Cells were equilibrated at 20°C for 3-4 hours, then spun at 50,000 rpm at 20°C, and scans collected every 30 seconds for 980 total frames. Scans were processed using DCDT+ to generate distributions of apparent sedimentation coefficients (g(s*))(75). An equivalent subset of 100 scans from each cell was included in fitting, corresponding to when boundary curves crossed the middle of the cell.

## Data Availability

All code used for structure processing and analysis is available on GitHub (https://github.com/kavranlab/KinaseTAPDimerScreen). Scripts are written in Python with the ProDy module for structure processing and analysis.

## Data Visualization

Kinome tree figures were generated using the CORAL webserver (76). Structures figures were generated in ChimeraX (77). Data were plotted using GraphPad Prism version 10.0.0 for MacOS, GraphPad Software, Boston, Massachusetts USA, www.graphpad.com.

## Acknowledgments

This work was supported by National Institutes of Health (NIH) Grant R35 GM156573 (to JMK). SB was supported by NIH T32 GM008403 and KAW by NIH T32CA009110. RG is supported by the NIH R37 GM059425, NIH R35 GM156244, and the Howard Hughes Medical Institute (HHMI). VLH was supported by NIH T32 GM007445 and the National Science Foundation (NSF) Graduate Research Fellowship Program (DGE2139757).

This manuscript is the result of funding in whole or in part by the NIH. It is subject to the NIH Public Access Policy. Through acceptance of this federal funding, NIH has been given a right to make this manuscript publicly available in PubMed Central upon the Official Date of Publication, as defined by NIH.

Computational work was carried out using the Advanced Research Computing at Hopkins (ARCH) core facility (rockfish.jhu.edu), which is supported by the NSF grant number OAC1920103. Molecular graphics and analyses performed with UCSF ChimeraX, developed by the Resource for Biocomputing, Visualization, and Informatics at the University of California, San Francisco, with support from NIH R01-GM129325 and the Office of Cyber Infrastructure and Computational Biology, National Institute of Allergy and Infectious Diseases. CD and AUC experiments were carried out at the Johns Hopkins University Center for Molecular Biophysics.

We would like to thank Laura Cordoba and Catherine Valadez for technical assistance.

