## supplemental materials for "A conserved dimerization element is required for protein kinase activation by *trans*-autophosphorylation"

**Supporting Information 1.** Crystal forms used in analysis.

Data is contained in an excel spreadsheet. The “All" tab contains each PDBID solved by X-ray crystallography of a human non-atypical eukaryotic protein kinase. The "Unique Forms" tab contains the PDBIDs used to represent each unique form. Representative crystal forms (same kinase, space group, and unit cell dimensions) are indicated in first column of the “All” tab.

**Supporting Information 2.** List of all kinase pairs and their BSA and pathway measurements

Data is contained in an excel spreadsheet. The “all pairs” tab contains each pair of kinase domains generated by applying crystal lattice symmetry to the asymmetric unit. Pair identifiers are reported using the format PDBID-chainID -chainID_SymOp, where SymOp is the PyMol symmetry operation code. The "reach" column indicates which pairs satisfy the AL modeling. Reported distance of AL lengths are equal to 3.4Å per residues in the AL. Rows shaded in gray in “all pairs” indicate trans-autophosphorylation compatible dimers and are replicated as a single set in the "t-ap compatible dimers" tab. Rows shaded in green in that tab correspond to the previously-documented complexes in **Figure 4A**.

**Figures**

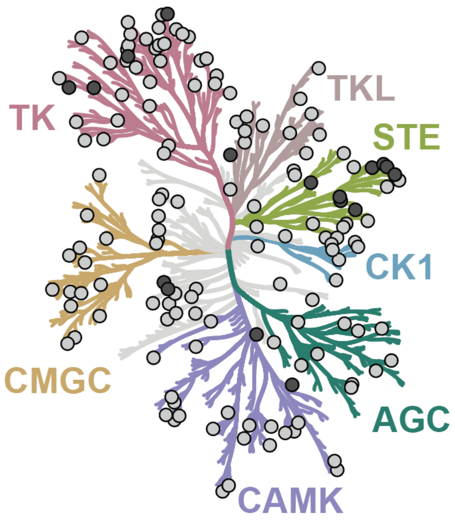


**Fig. S1.** *Distribution of trans-autophosphorylation dimers.* Human kinome tree with dark gray spots indicating kinases with previously identified structures of *trans*-autophosphorylation dimers and light gray dots indicating additional dimers compatible with *trans*-autophosphorylation identified in this study.

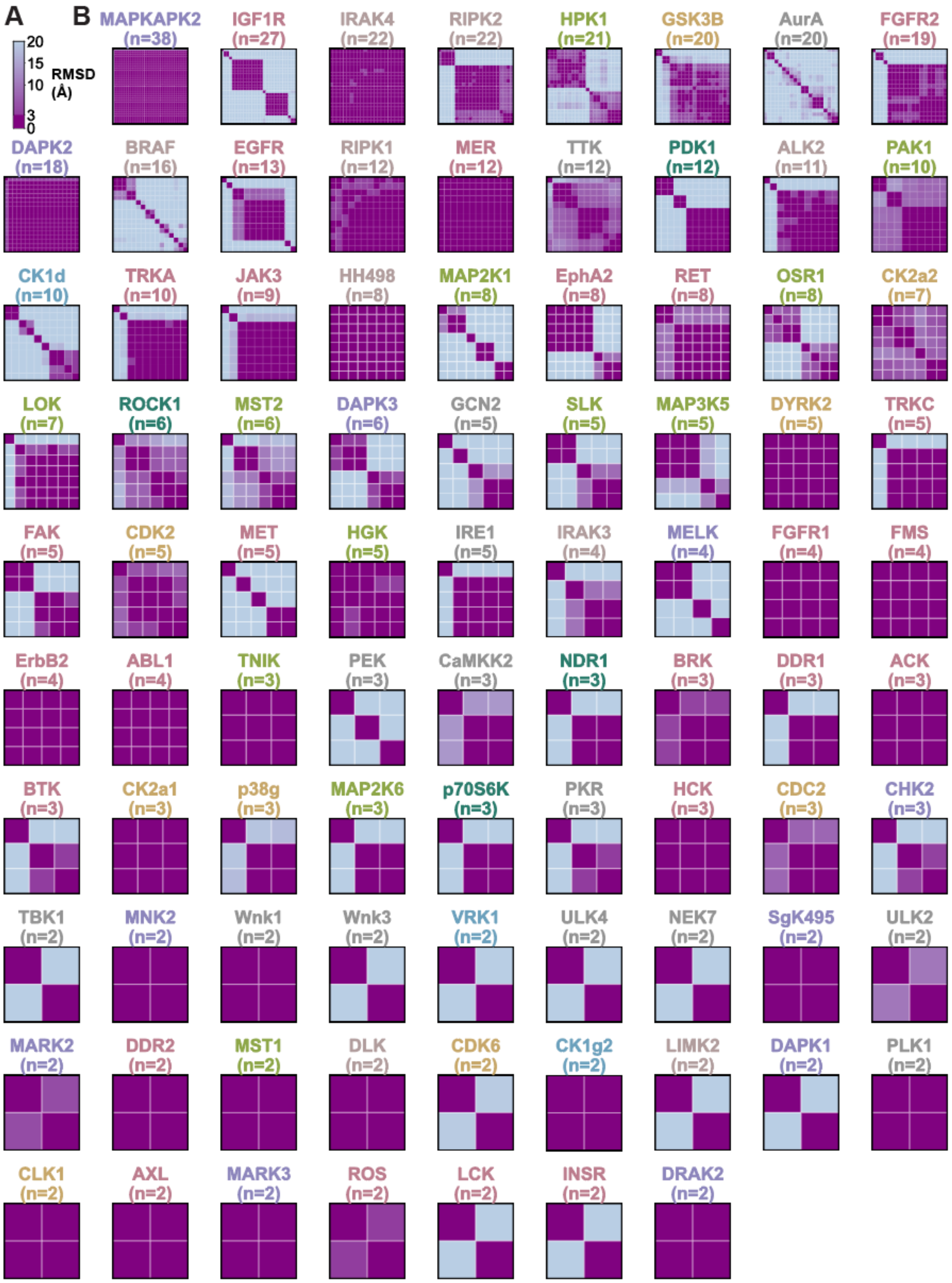


**Fig. S2.** *Pairwise superpositions of individual kinase with 2 or more trans-autophosphorylation compatible dimers.* (A) RMSD color key. (B) Heatmaps showing pairwise superpositions of all structures for each kinase with two or more *trans*-autophosphorylation compatible dimers, ordered from most to fewest observations.

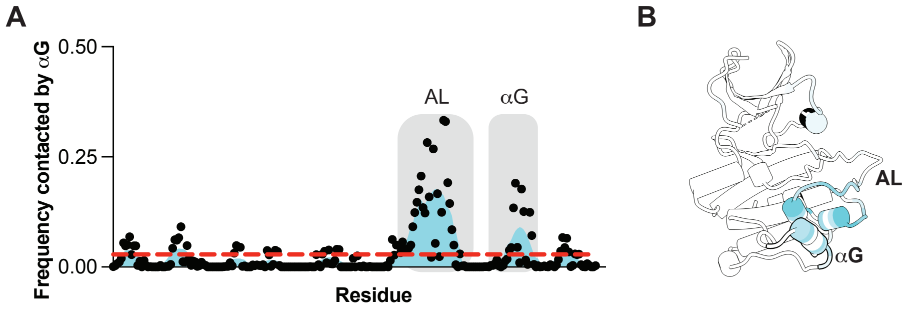


**Figure S3.** *αG preferentially binds itself and the AL on the partner kinase.* (A) Frequency with which residues on the partner kinase are contacted by αG in αG-mediated dimers. The mean frequency is indicated by a dashed red line, and the frequency was fit to a smoothed function (cyan) for visualization purposes. The two most frequently buried regions are indicated by gray boxes. (B) Cartoon representation of phosphorylated MST1 kinase domain (PDBID 3COM) colored by frequency contacted (dark cyan high, light cyan low).

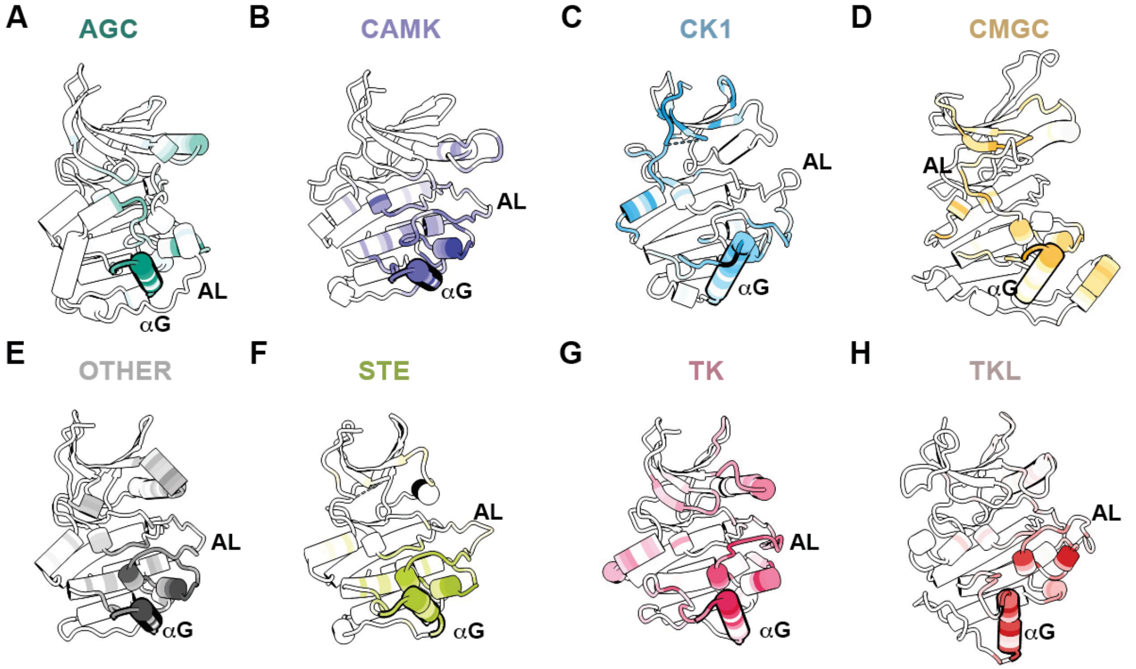


**Figure S4.** *Frequency residues are buried in trans-autophosphorylation compatible dimers by kinome group.* Cartoon of a representative kinase from each kinome group colored by the frequency residues are buried at the interface (dark, high frequency; light, low frequency). Kinases shown are: (A) NDR1 (6BXI), (B) DAPK3 (1YRP), (C) CK1γ2 (2C47), (D) JNK2 (3NPC), (E) AuroraA (3E5A), (F) MST1 (3COM), (G) FES (3BKB), and (H) IRAK4 (2NRU).

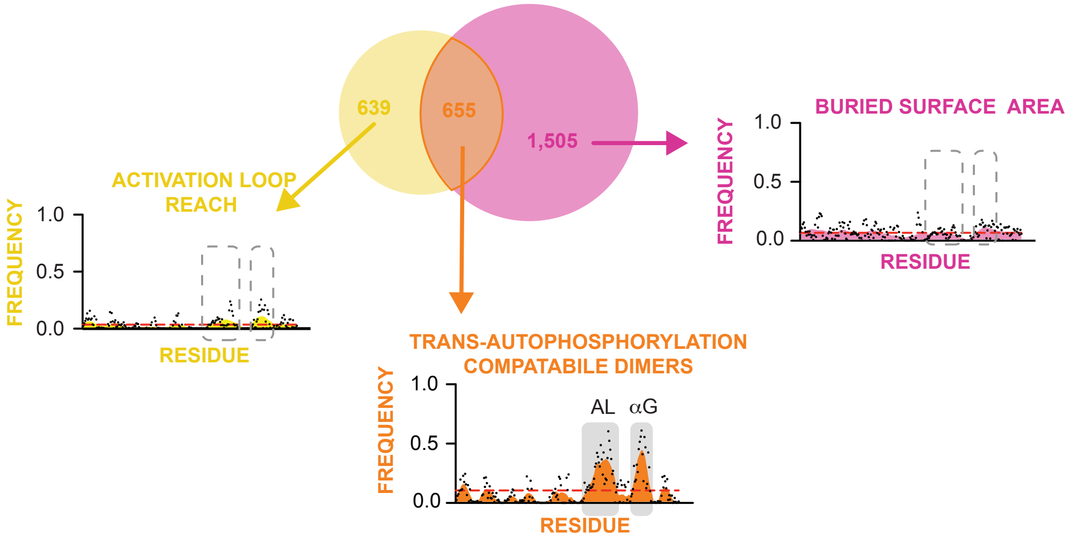


**Figure S5.** *Comparison of computational screen criteria and their contribution to αG enrichment.* Venn diagram showing the number of kinase domain pairs satisfying each criterion either independently (yellow or pink) or both simultaneously (orange). Dimer footprint plots show the frequency with which residues in conserved secondary structure elements are at the dimer interface for each subset. Dashed red line indicates mean frequency; smoothed fit (yellow, pink, or orange) shown for visualization.

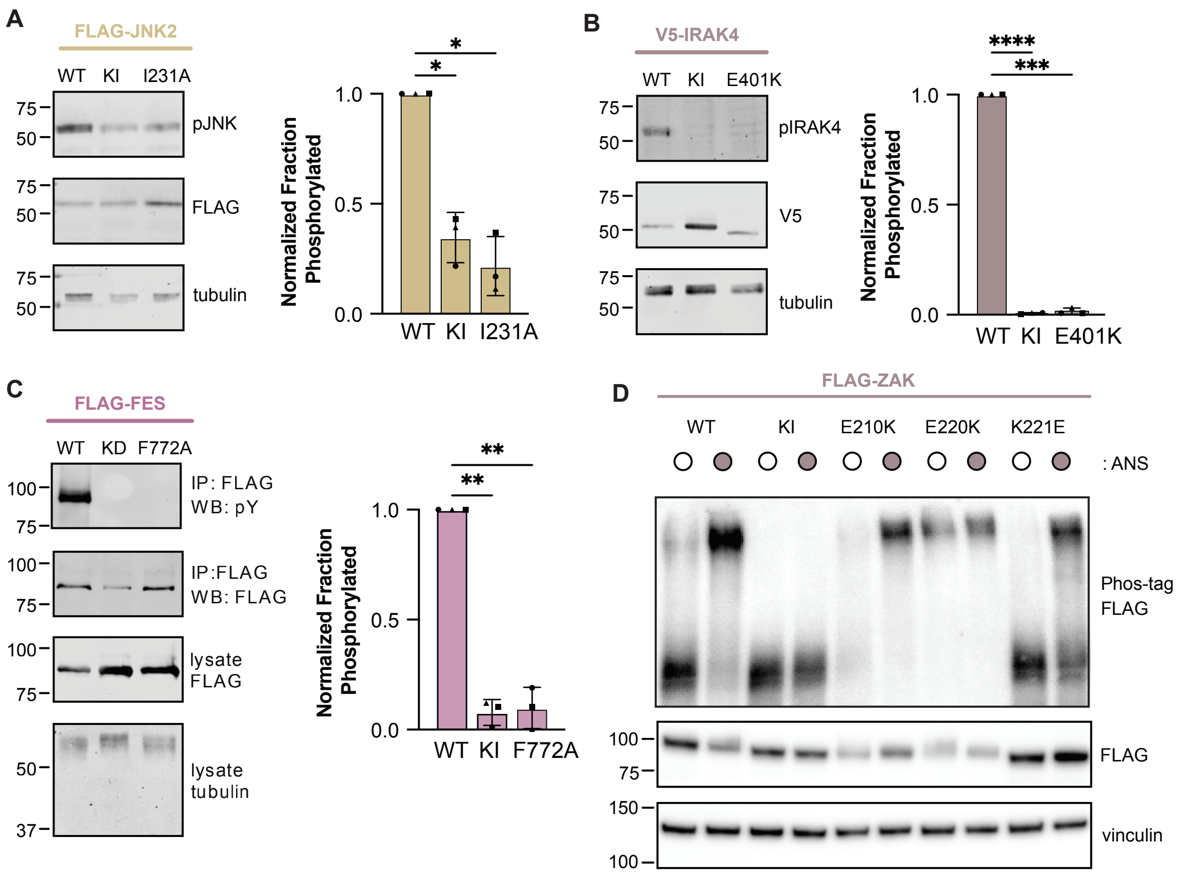


**Figure S6.** *Single-residue substitutions of $\alpha$G disrupt AL autophosphorylation of full-length kinases in cells.* (A,B) Representative set of Western blots of HEK293 lysates transiently transfected with indicated variants of JNK2 or IRAK4, respectively, and probed with indicated antibodies. Bar graphs of the mean with standard deviation of fraction phosphorylated protein normalized to wild-type for each protein. Biological replicates are indicated by different shaped points. Significance determined by one-way ANOVA followed by Dunnett's test. P values used: ****<0.0001; *** <0.001, ** < 0.01; *<0.05, and 0.05> ns. (C) Same as panel B but for FES isolated by immunoprecipitation. (D) HEK293 ZAK knockout cells transiently transfected with FLAG-ZAK variants, as indicated. ZAK phosphorylation and expression monitored by Phos-tag in the presence or absence of anisomycin.

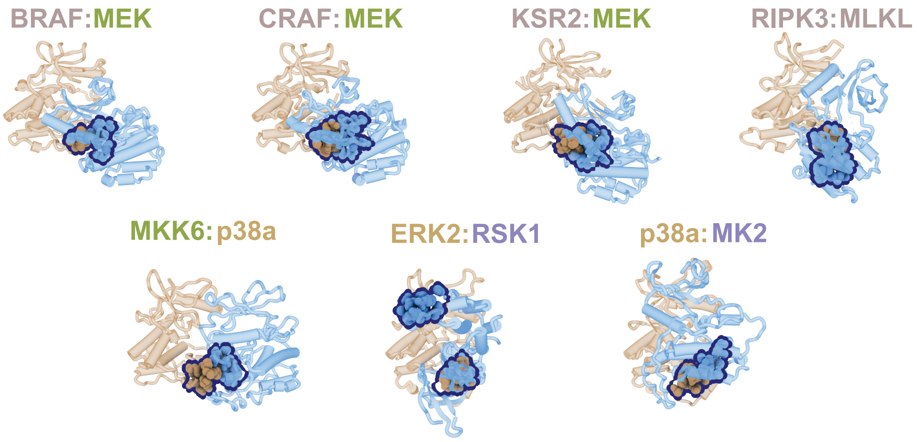


**Figure S7.** *αG mediates dimerization during trans-phosphorylation.* Cartoon representation of *trans*-phosphorylation dimers shown with the upstream kinase (*e.g.* the enzyme) in tan and the downstream kinase (*e.g.* the substrate) in light blue. αG residues are displayed in spheres and outlined. PDBIDs of structures displayed are: *Row 1* – 6Q0J, 8CHF, 2Y4I, and 7MON*; Row 2* -- 8A8M, 4NIF, and 6TCA.

**Tables**

**Table S1.** Kinase Variants Used

| Kinase | Variant | Substitution(s) |
|---|---|---|
| CHK2 | KI | D368A |
| CHK2 | αG* | R431A, T432A, Q433A, V434A, S435A, K437A, D438A, T441A, S442A |
| FES | KI | K590R |
| FES | αG* | Q768A, F772A, K775A |
| IGF1R | KI | D1135A |
| IRAK4 | KI | D329A |
| IRAK4 | αG* | Q394A, L395A, D398A, E401A, E404A, D405A, E406A |
| JNK2 | KI | K55R |
| JNK2 | αG* | T228A, D229A, H230A, I231A, E239A |
| NDR1 | KI | K118A |
| NDR1 | αG* | E325A, T326A, Q328A, K332A, M335A, N336A |
| ZAK | KI | K45M |
| ZAK | αG* | E210K, L212A, W216A, E220K, K221E |

**Table S2.** Source and lots of antibodies used

| **Antibody** | **Manufacturer** | **Lot #** |
|---|---|---|
| V5-Tag (D3H8Q) Rabbit | Cell Signaling Technology | 7 (CHK2,NDR1)<br>8 (IRAK4) |
| FLAG M2 Mouse (for FES & JNK2) | Millipore Sigma | 1003602111 |
| FLAG M2-HRP Mouse (for ZAK) | Millipore Sigma | 0000480821 |
| IGF-1Rβ Rabbit | Cell Signaling Technology | 4 |
| Phospho-CHK2 (T383) | Abcam | 1001308-1 |
| Phospho-SAPK/JNK (T183/Y185) (98F2) Rabbit | Cell Signaling Technology | 10 |
| Phospho-Tyrosine (4G10) Mouse | Cell Signaling Technology | 1 |
| Phospho-IGF-1Rβ (Y1135) Rabbit | Cell Signaling Technology | 2 |
| Phospho-IRAK4 (T345/S346) (D6D7) Rabbit | Cell Signaling Technology | 8 |
| Vinculin | Santa Cruz | H3023 |
| Tubulin | Cell Signaling Technology | 7 |
| IRDye® 800CW Goat anti-Mouse IgG | LICORbio | N/A |
| IRDye® 800CW Goat anti-Rabbit IgG | LICORbio | N/A |
| HRP-anti-mouse (secondary) | Cell Signaling Technology | 40 |
| HRP-anti-rabbit (secondary) | Cell Signaling Technology | 34 |
